# Widespread cortical representations of innate behaviors in the mouse

**DOI:** 10.1101/2025.06.05.657711

**Authors:** Nicholas J. Michelson, Pankaj K. Gupta, Timothy H. Murphy

**Author notes:** Correspondence to: Timothy H Murphy, Address: 2215 Wesbrook Mall, Vancouver, BC, V6T 1Z3, Canada.

## Abstract

Across cortex, a substantial component of observed neuronal activity can be explained by movement. Voluntary movements elicit cortical activity that contain elements related to both motor planning and sensory activity. However, automatic movements that do not require conscious processing, for example grooming or drinking, are to a large extent controlled by subcortical systems such as the brainstem. While the cortex may not be required to generate these automatic behaviors, it is unclear whether cortical activity represents the movements associated with them. In this work, we use a simple procedure to stimulate stereotyped grooming behaviors in the head-fixed mouse while measuring neuronal function across the dorsal cortex. We find specific cortical representations of grooming both at a mesoscale network-level and single-cell resolution. Mesoscale cortical activation was most prominent at the onset of grooming episodes, and declined to baseline levels despite continuous engagement in the behavior. Further stratification of grooming component movements revealed that more directed and unilateral grooming movements had greater cortical associated responses than more stereotyped bilateral movements. These findings help to frame the impact of forms of automatic movements on large scale cortical ensembles and suggest they engage specific and transient cellular and regional cortical ensembles.

## Introduction

Movement elicits widespread neuronal activation across the cortex and can affect other cortical functions. For example, locomotion increases neuronal firing rate responses to stimuli in the visual cortex without affecting orientation tuning properties (Niell and Stryker 2010). Locomotion also desynchronizes mesoscale cortical networks (West, Gerhart, and Ebner 2024) while enhancing correlations between single neurons and distal cortical areas (Clancy, Orsolic, and Mrsic-Flogel 2019). Besides locomotion, widespread cortical activation is observed during spontaneous movements of the paws and face (Musall et al. 2019; Stringer et al. 2019). Cortical neuronal ensembles therefore encode mixed representations of behavior and sensory or task information (Stringer et al. 2019; Vinck et al. 2015; Musall et al. 2019). Investigating cortical representations of movement during different behavioral contexts may therefore help to elaborate the general principles by which cortical neurons integrate motor, sensory, and cognitive information.

Voluntary movements, while controversial in their definition (Prochazka et al. 2000), are generally considered to be cortex-dependent actions that contain elements related to motor planning, movement, and sensory activity (Tibbetts 2013). In the mouse, optical inhibition of sensorimotor cortical areas impairs voluntary forelimb movements, such as reaching for a food pellet or water drop (Galiñanes, Bonardi, and Huber 2018; Guo et al. 2015; Yang, Kanodia, and Arber 2023). These cortical motor areas send topographically organized input to the brainstem which facilitates specific phases of the movement, e.g. reaching for the food pellet or handling the pellet during consumption (Yang, Kanodia, and Arber 2023). However, while cortical inhibition impairs skilled motor behaviors, the execution of untrained behaviors such as grooming (Guo et al. 2015), is unaffected, despite apparent similarities in the movement (Naghizadeh, Mohajerani, and Whishaw 2020). Moreover, grooming behaviors can be expressed in decerebrate rodents (Berntson, Jang, and Ronca 1988; Berntson and Micco 1976) and induced by stimulation of the lateral rostral medulla (Ruder et al. 2021), indicating that the neural substrate for generating this behavior exists within the brainstem. Thus, skilled forelimb behaviors are facilitated through communication between the motor cortex and the brainstem, while innate behaviors such as grooming or drinking can be expressed to some degree with only the brainstem circuitry.

Is the widespread cortical representation of movement observed during subcortically mediated behaviors? Multiphoton calcium imaging experiments in the forelimb motor cortex revealed functionally and spatially segregated neuronal ensembles during running and grooming behaviors (Dombeck, Graziano, and Tank 2009) and extracellular electrophysiological recordings in the forelimb motor cortex have shown that spike rate changes occur during phase transitions within grooming sequences (Sjöbom et al. 2020). Thus, cortical correlates of subcortical behaviors have been documented. However, the spatial scale of investigations is limited by recording methodologies. In this work, we expand upon previous efforts to characterize cortical network activity during grooming. Neuronal activity across the dorsal cortex is assessed using single-photon mesoscale cortical calcium imaging and wide field-of-view two-photon calcium imaging in head-restrained mice while grooming behaviors are evoked with a water drop placed onto the orofacial area. This stimulus reliably evokes grooming behaviors, which exhibit a mix of the stereotyped movements observed during naturalistic grooming (Aldridge and Berridge 1998) and movements that resemble reaching behaviors directed towards the stimulus (Naghizadeh, Mohajerani, and Whishaw 2020). Distinct mesoscale cortical activation patterns were observed across different grooming component behaviors, and large cortical responses were observed at the onset of grooming movements, which subside over extended periods of continuous behavior. This decline in cortical activity is also observed during extended periods of continuous licking and drinking behaviors. At the single-cell level, a distributed neuronal population across cortical areas exhibited activity that is significantly more correlated with grooming than non-grooming movements. Taken together, our results suggest that the cortex reflects subcortically-mediated behaviors through a large transient response at the onset of the behavior, and a sustained response across a distributed population of neurons.

## Materials and Methods

### Animals

All procedures were approved by The University of British Columbia Animal Care Committee and conformed to the Canadian Council of Animal Care and Use guidelines (protocol A22-0054). Single-photon imaging experiments were performed on transgenic mice (n=4, 3 to 4 months old) expressing GCaMP6s (Chen et al. 2013). Three transgenic lines were used in this study with the goal of imaging calcium activity in excitatory neurons: 1) Thy1-GCaMP6s (n=1 female; Jackson mouse #024275, GP4.3, C57BL/6-Tg(Thy1-GCaMP6s)GP4.3Dkim/J16)(Dana et al. 2014); 2) CAMKIIa-GCaMP6s (n=2 females; crossing of two lines: #024742, B6; DBA-Tg(tetO-GCaMP6s)2Niell/J, B6 with #003010 B6;CBATg(Camk2a-tTA)1Mmay/J)(Wekselblatt et al. 2016); and 3) TIGRE-GCaMP6s (n = 1 male) (crossing of three lines: #005628 B6.129S2-Emx1^tm1(cre)Krj^✉J and #007004 B6.Cg-Tg(Camk2a-tTA)1Mmay/DboJ and #024104; Ai94; B6:Cg-Igs7^tm94.1(tetO-GCaMP6s)Hze^✉J strains (Madisen et al. 2015). Multi-photon imaging experiments were performed on transgenic mice expressing GCaMP6s (Thy1-GCaMP6s (Dana et al. 2014), n=1 male and n=1 female; and Camk2a-tTA x tetO-GCaMP6s (Wekselblatt et al. 2016), n=1 female) or jGCaMP8s (Camk2a-tTA x tetO-jGCaMP8s, n=1 male).

### Surgical procedures

Animals were anesthetized with 3% isoflurane and maintained at 1-2.5% over the course of the procedure. After induction, ophthalmic ointment was applied to both eyes and body temperature was maintained at 37.1C using a closed-loop rectal temperature regulator. The fur on the top of the head was removed using surgical scissors, and the skin was cleaned using a triple scrub of 0.1% betadine in water followed by 70% ethanol. Local analgesic was then applied through an injection of lidocaine (0.1 ml, 0.2%) under the scalp. The mice were also administered subcutaneously with a saline solution containing meloxicam (2⍰⍰mg/ml), atropine (3μg/ml), and glucose (20 mM). The skin on the top of the head was then resected, and fascia and connective tissues on the surface of the skull were removed so that the skull surface was completely clear of debris and dry.

For single-photon widefield imaging, a titanium bar (22 × 3.25 × 2.8 mm) was fixed to the cerebellar plate, posterior to lambda using Krazy glue, and then reinforced with clear dental cement. Clear dental cement was prepared by mixing 1 scoop of Metabond powder with 7 drops of C&B Metabond Quick Base, and 1 drop of C&B Universal catalyst (Parkell, Edgewood, New York). A layer of clear dental adhesive was also applied onto the skull, and a precut cover glass was placed gently on top of the mixture before it solidified.

For two-photon imaging, a triple-layer glass window was prepared prior to surgery by attaching two 5mm diameter circular glass coverslips (Warner Instruments) with ultraviolet (UV) curing glue (Norland products #6101), which were then glued to a larger 6mm diameter circular glass coverslip, resulting in a triple layer glass window with a plug (Hattori and Komiyama 2022). UV light was applied with a UV flashlight (Alonefire SV43). A 5mm diameter circular craniotomy was performed using an air-powered dental drill and the triple layer glass window was placed into the craniotomy with the glass plug gently pressing on the surface of the brain. The window was first fixed in place using Krazy glue, and a titanium headbar was placed over lambda. The entire preparation including the edges of the cranial window and the titanium headbar, was reinforced with darkened dental cement (Araragi, Alenina, and Bader 2022). Dental cement was darkened by mixing carbon powder (Sigma-Aldrich, 484164) into the dental cement.

### Behavior imaging

Mice were head-restrained in a plexiglass apparatus and the scene was illuminated with 850nm LEDs (Gupta and Murphy 2025). A monochrome camera (Omron-Sentech, STC-MBS43U3V) connected to a NVIDIA Jetson was equipped with a bandpass infrared filter (840-865 nm, Bock Optronics) and positioned directly in front of the animal to record 8-bit behavior videos at 90 frames per second with a resolution of 640 × 320 pixels (Figure 1A). After an initial baseline period, grooming behavior was evoked by delivering a drop of water onto the orofacial area once per minute for 20 minutes. Each water drop stimulus was preceded by a 1-second 10kHz tone followed by a 1-second delay period (Figure 1B). For each mouse, at least two spontaneous sessions, which contained the audio cue without the water drop stimulus, were recorded before the evoked grooming sessions (Figure 1C).

**Figure 1.**
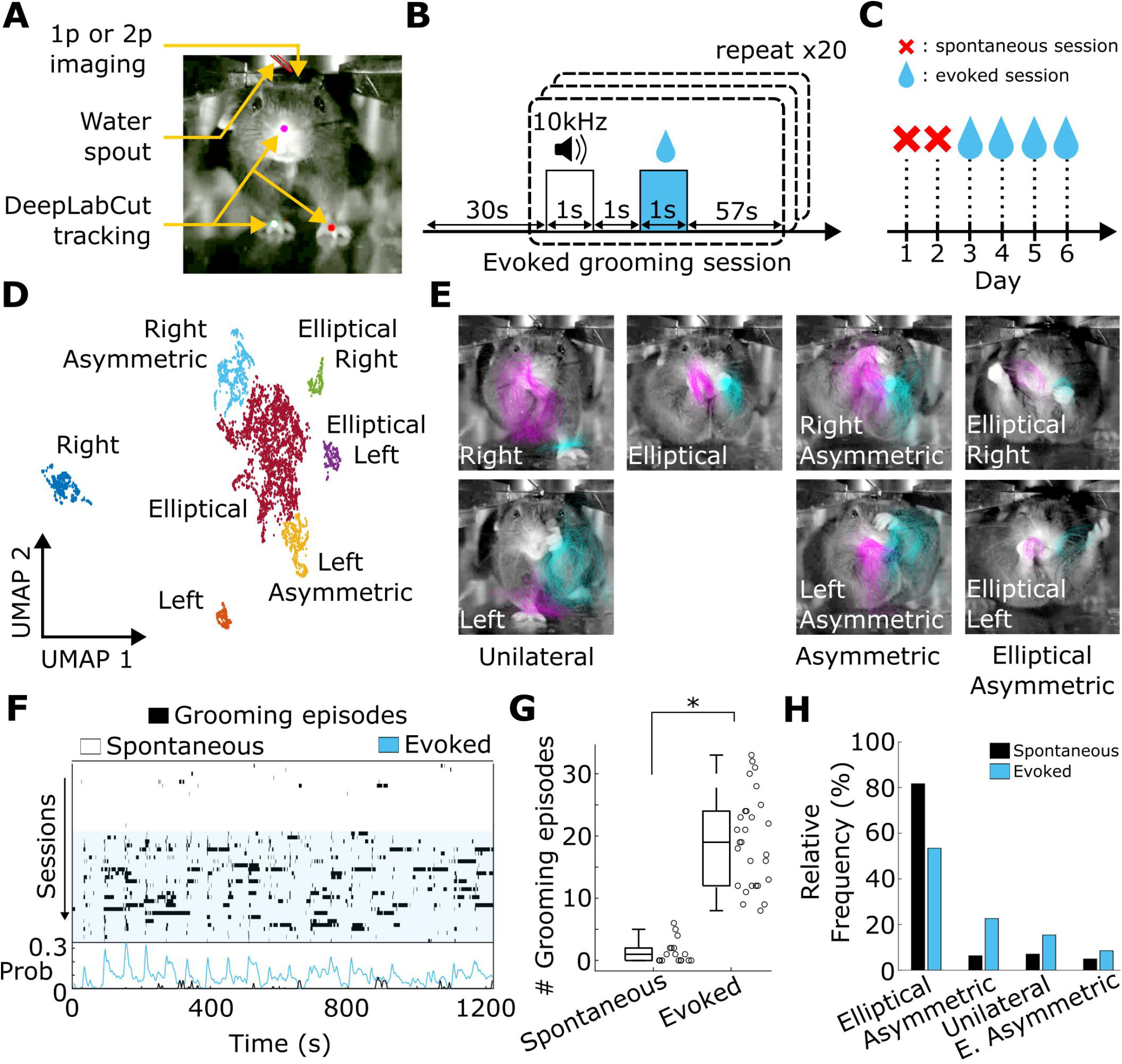
Water drop stimulus paradigm evokes grooming behaviors with a rich diversity of movements. **A)** Experimental setup. **B)** Trial structure for an evoked grooming session. **C)** Experiment timeline. **D)** Visualization of paw trajectories in UMAP space. Each point represents an individual behavior. Points are colored by cluster. **E)** Example behaviors from clusters in (D). Colored lines show paw trajectories for 200 individual example behaviors in each behavior cluster. **F)** Rasters of grooming events during spontaneous sessions without a water drop stimulus (white), and evoked sessions with 20 water drop stimuli (blue). Probability of grooming (bottom) for spontaneous and evoked sessions. **G)** Quantification of number of grooming episodes. p=2.1e-12, t-test. **H)** Relative frequency of each grooming behavior during spontaneous and evoked sessions. Images in (A) and (E) are adjusted for brightness to enhance visibility.

### Identification of grooming behaviors

Grooming behaviors were identified using two independent approaches - manual labelling (Figure 1-figure supplement 1A) and unsupervised clustering (Figure 1D). Final results are presented using the unsupervised clustering approach, however both labelling strategies demonstrated substantial agreement (Figure 1D, Figure 1-figure supplement 1B) and the results presented throughout the paper were consistent across approaches. For manual labelling, the start and end index for each grooming event were saved and manually curated using Behavioral Observation Research Interactive Software (BORIS) (Friard and Gamba 2016). Grooming component behaviors were classified by visual inspection as one of seven distinct behaviors, including unilateral facial strokes: right and left; and bilateral facial strokes: elliptical, elliptical asymmetric, right asymmetric, left asymmetric, and large bilateral (Figure 1- figure supplement 1B). BORIS was also used to label licking behaviors where the tongue clearly protruded from the mouth, as well as the precise timepoints in which the full weight of the drop makes contact with the mouse.

For unsupervised labelling, the position of the paws and nose were tracked using DeepLabCut (Mathis et al. 2018). A single point was used for each paw and around 20-60 frames from each trial and mouse were selected for labeling. The network was trained with default parameters for 1030000 iterations. The start and end frame index of each grooming behavior obtained using BORIS were used to extract paw position and velocity information during each behavior event. This information was distilled down to 44 features for each behavior event (Table 1), yielding a matrix of size Nx44, where N is the number of grooming behaviors exhibited across all mice. This feature matrix was then reduced to 2 dimensions using Uniform Manifold Approximation and Projection (UMAP) for dimension reduction. The resulting embedding exhibited several spatially distinct clusters, which were extracted with hierarchical clustering (Figure 1D). These clusters correspond with distinct and stereotyped patterns of paw movements (Figure 1E, Figure 1-figure supplement 1C).

### Quantification of grooming behaviors

To assess the sequential patterning of behavior within grooming episodes, component behaviors were aggregated such that any grooming behavior occurring within a 3-second window from another would be grouped into the same bout. The duration of the aggregation window (3-s) was chosen empirically, however selecting aggregation window durations ranging from 1-s to 10-s did not substantially affect the results. Licking events often overlap with grooming behaviors and therefore were not considered for this analysis. A transition matrix, tracking the number of transitions between behaviors, was created for each grooming episode and the resulting transition matrices were then summed across episodes within a single session and then averaged across mice and sessions (Figure 2D). The organizational structure of grooming episodes was then assessed using Louvain community detection (Rubinov and Sporns 2011), which partitions the behavior transition matrix into clusters which maximize the overall modularity of the network. The resulting communities represent groups of behaviors with frequent transitions within groups and infrequent transitions between groups.

**Figure 2.**
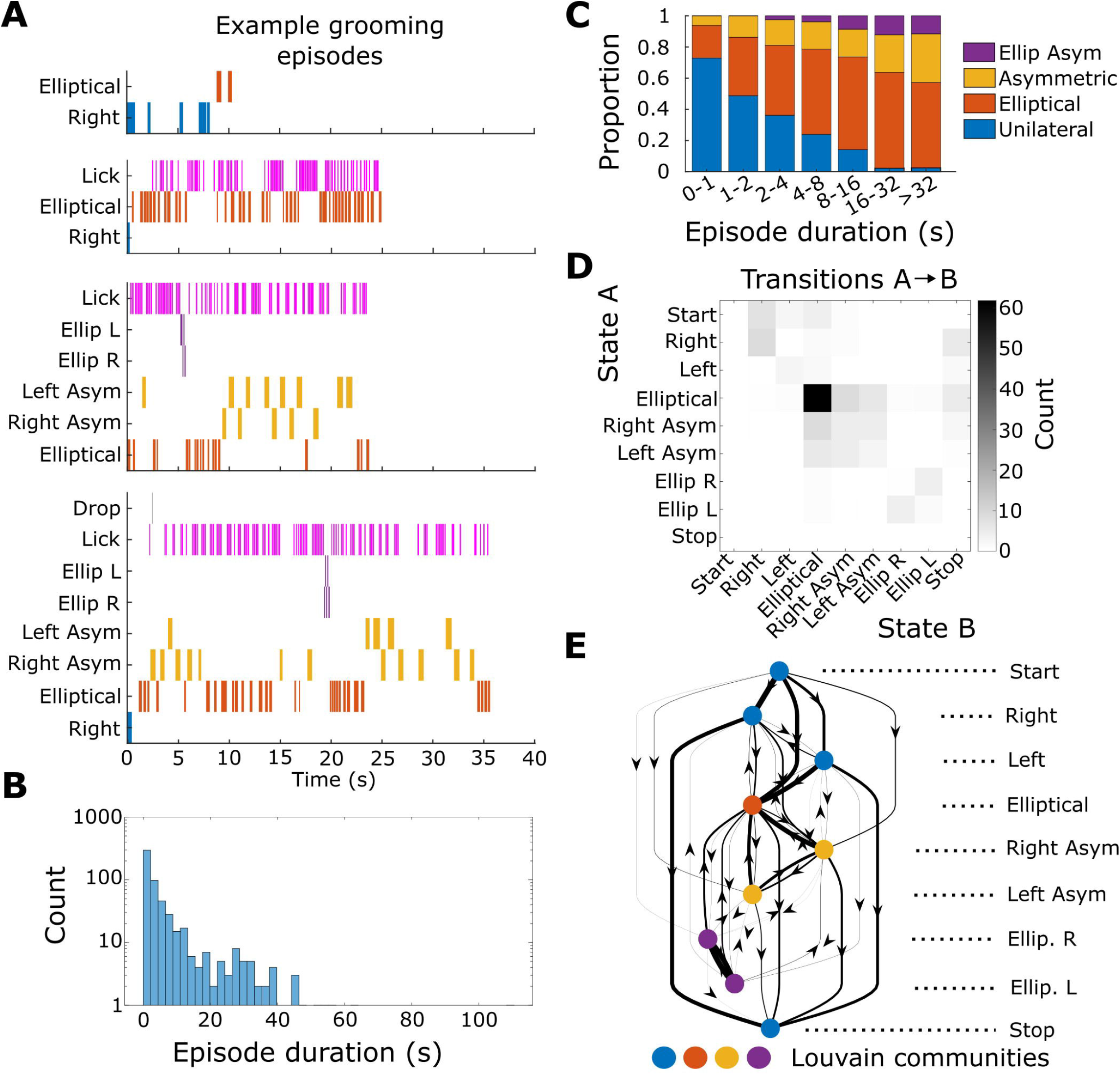
Temporal presentation of grooming behaviors. **A)** Example grooming episodes. Note that the examples shown here are longer than the average grooming events (see panel B). These examples are intended to showcase the variability in constitution of grooming episodes. **B)** Histogram of episode duration. **C)** Proportional composition of grooming episodes of various durations. Data compiled across all mice. **D)** Average transitions between grooming behaviors across all mice. **E)** Directed network representation of transition matrix in (D). Nodes are arranged using MATLAB’s ‘layered layout’ organization. Edge thickness is proportional to transition probability between nodes. Self-connections are omitted for clarity. Nodes are colored by Louvain community assignment performed on the transition matrix.

### 1-photon imaging

Dorsal cortical GCaMP activity was recorded using similar methods as described previously (Ramandi et al. 2023). Briefly, images of the cortical surface (Figure 3A) were projected through a pair of back-to-back photographic lenses (50 mm, 1.4 f:135 mm, 2.8 f or 50 mm, 1.4 f:30 mm, 2 f) onto a 1M60 Pantera CCD camera (Dalsa). GCaMP was excited with a blue LED (Luxeon, 473 nm) with a band-pass filter (Chroma, 467–499 nm) delivered to the surface of the cortex through a liquid light guide (Thorlabs). GCaMP fluorescence emission was filtered using a 510–550 nm band-pass filter (Chroma). 12-bit images were collected at 30 frames per second using XCAP imaging software using 8×8 pixel on-chip binning, yielding images of size 128×128 pixels. Video acquisition was triggered by the Jetson which also started behavior video recording, and after a few seconds of delay, the blue cortical excitation LED and the infrared behavior LED were simultaneously turned on by TTL from the Jetson. At the end of the trial, the cortical and behavior LEDs were turned off simultaneously prior to stopping acquisition. Frames were synchronized across behavior and brain cameras by matching the illuminated frames at the start and end of the trial.

**Figure 3.**
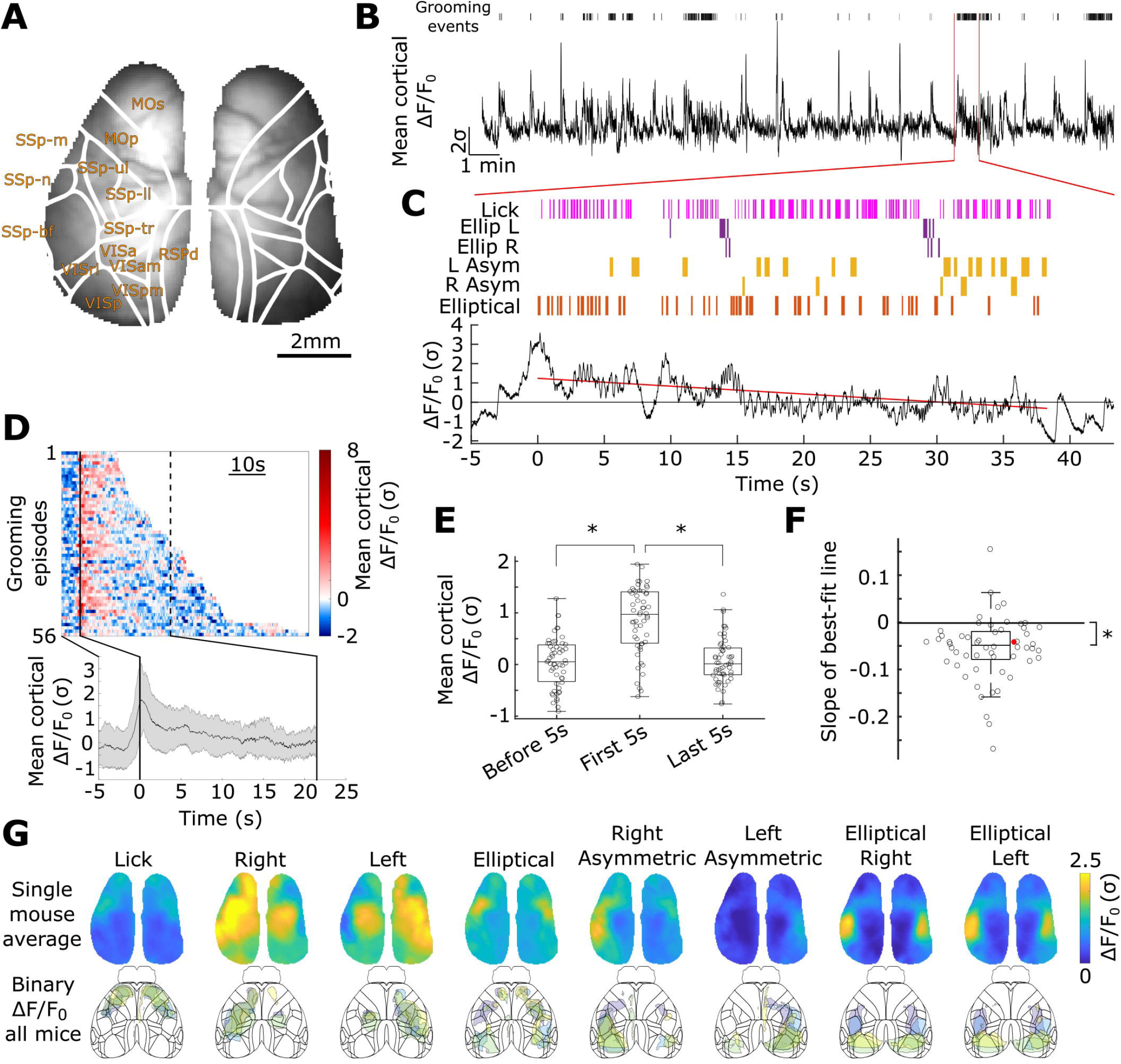
Mesoscale cortical widefield activation during grooming. **A)** Example image of GCaMP6s fluorescence in a Thy1-GCaMP6s mouse with the Allen Institute cortical atlas overlaid. (Abbreviations - MOs: secondary motor area; MOp: primary motor area; SSp-m: primary mouth somatosensory area; SSp-n: primary nose somatosensory area; SSp-ul: primary upper limb somatosensory area; SSp-ll: primary lower limb somatosensory area; SSp-tr: primary trunk somatosensory area; SSp-bf: primary barrel field somatosensory area; VISa: anterior visual area; VISrl: rostrolateral visual area; VISam: anteromedial visual area; VISpm: posteromedial visual area; VISp: primary visual area; RSPd: dorsal retrosplenial area) **B)** Averaged activity dF/Fo across the entire cortical surface. Grooming behaviors indicated at the top (black bars). **C)** Ethogram (top) and mean cortical calcium activity (bottom) during the grooming episode indicated in (B). Best fit line through cortical activity shown in red. **D)** Heatmap (top) of averaged activity across the entire cortical surface for all grooming episodes which exceeded 10s. The dotted line at ∼21s is drawn at the 50th percentile duration. The mean cortical response over rows in the heatmap. Shading shows the standard deviation. **E)** Comparison between averaged cortical signals at various time points. Lines indicate significance. p<0.05 repeated measures ANOVA with post-hoc Bonferroni correction. **F)** Slope of the best fit line for all grooming episodes exceeding 10s. Slope is less than 0. p<0.05 t-test. Red marker indicates example shown in (C). **G)** Response maps for a single mouse averaged across all frames in which the behavior is expressed (top). Averaged response maps for each mouse binarized at >80th percentile to show maximally activated networks (bottom). Each colored contour reflects a different mouse.

### Drinking experiments

Drinking experiments were conducted after all grooming experiments were completed. Data acquisition and analysis were similar as with other 1-photon imaging data, except that behavior videos were acquired at 30 frames per second. Mice were water-limited by removing the water bottles from the homecage. During the experiment, water was provided to the mouse via a water spout placed directly in front of the mouth. The delivery of the water was controlled in the same way as in the grooming experiments. After imaging, mice were provided with supplemental access to water for ∼15-30 minutes to ensure adequate daily intake. The weights of the mice were monitored daily to ensure weight loss did not exceed 15% of the initial weight (Gupta and Murphy 2025).

### 1-photon image pre-processing

Single photon wide-field fluorescence data was compressed using singular value decomposition (SVD), which yielded *U*, the matrix of pixels × components; *V*, the matrix of components x time; and s the diagonal matrix of singular values. The top 1,000 components were retained, and s was multiplied into V, which was then upsampled to 90 fps to match the behavior video. The matrix *s*\**V* was then high-pass filtered at 0.01 Hz using a second-order zero-phase Butterworth filter. Wide-field fluorescence data were combined within mice across separate recordings, by recasting experiment-specific SVD components into a master SVD basis set (Peters et al. 2021). To create the master SVD basis set, spatial components from each imaging session were first aligned to a reference session and concatenated, then SVD was performed on the resulting matrix and the top 1,000 components were retained. Temporal components for each experiment then were recast into the master basis set as described in (Peters et al. 2021). Wide-field fluorescence data for each mouse were then aligned to the Allen Institute Common Coordinate Framework (Q. Wang et al. 2020) using a control-point based registration method with key points placed along the superior sagittal sinus, bregma, and the space between olfactory bulbs and frontal cortex (Saxena et al. 2020). A mask was drawn manually over the cortex of the reference image for each mouse to discard pixels outside of the cortex from analysis.

### photon image analysis

Fluorescence activity was quantified by calculating the z-scored DF/F0. First, imaging data were reconstructed from the spatial and filtered temporal SVD components. Since temporal filtering removes the DC offset, the minimum value from the entire data matrix was subtracted from each pixel’s data, ensuring that the mean across time was positive for each pixel. The DF/F0 was then computed by dividing each pixel by its average intensity over the entire session, and then subtracting 1. Finally, the DF/F0 was z-scored across time. To create averaged network activation maps, cortical frames that were concurrent with the respective behavior were indexed and stored in a separate array. The average of these frames were taken across sessions (Figure 3G top). To assess maximally activated networks across behaviors, the 80th percentile DF/F0 value was computed for each mouse’s averaged cortical activation map, and used as a threshold to create a binary map for each mouse and behavior (Figure 3G bottom).

### Ridge regression

The linear model analysis was based off of the code provided by Musall et al (Musall et al. 2019). Briefly, the binary behavior vectors were used to create a design matrix which comprised separate columns of time-shifted pulses ranging from -0.5 to 2s from the onset of each movement and from 0 to 1s from the onset of each stimulus (water drop and audio cue). The design matrix was fit using ridge regression to the filtered and upsampled neural fluorescence SVD temporal components, resulting in time-varying coefficients for each behavior which can then be multiplied by the neural spatial SVD components to visualize each behavior’s estimated cortical responses (Figure 3-figure supplement 2B). The model fit was estimated by computing the explained variance using tenfold cross-validation (Figure 3-figure supplement 2C). To estimate a variable’s unique contribution to the data, a reduced model was created by randomly permuting the binary vector for that variable prior to creation of the design matrix, thereby eliminating any meaningful relationship between that behavior and the neural data. The difference in explained variance between the full model and the reduced model was then computed to assess that variable’s unique contribution to the data (Figure 3-figure supplement 2D).

### 2-photon imaging

Two-photon imaging was performed using the Diesel2p microscope (Yu et al. 2021) controlled with MATLAB software Scanimage (Vidrio). Two-photon laser excitation was provided by an 80-MHz Newport Spectra-Physics Mai Tai HP 1020. The Diesel2p’s dual scan engines were utilized to maximize the imaging area and imaging regions of interest (ROIs) were chosen across mice and imaging sessions to cover as many cortical regions as possible. Each scan path consisted of a resonant scanner (CRS 8 KHz, Cambridge Technologies), and two XY galvanometers (8320K, Cambridge Technologies). While imaging area sizes sometimes varied between scan engines, the spatial resolution was held constant across scan arms, resulting in acquisition rates that varied, with larger areas collected at lower frames per second. Photon signal was collected by photomultiplier tube (H11706P-40, Hamamatsu) and amplified with a high-speed current amplifier (HCA-400M-5K-C, Laser Components). Imaging was performed with 920 nm laser at ∼80 mW excitation out of the front of the air objective (0.55 NA). Behavior video acquisition and resonant scanning were initiated prior to collecting 2-photon images. Acquisition of 2-photon images was triggered using TTL output from the Jetson which simultaneously turned on infrared LEDs which illuminated the scene, and at the end of the trial, 2-photon image acquisition was terminated by TTL and the infrared LEDs were simultaneously turned off. Behavior video acquisition ended following a short delay after turning off the LEDs and 2-photon images were synchronized to the behavior video by matching to the illuminated frames.

### 2-photon image processing

2-photon images were spatially binned by a factor of 2×2 to improve signal to noise ratio and reduce file sizes for subsequent operations. After binning, neuron diameters were typically ∼5-8 pixels. Motion correction and neuron detection were then performed using Suite2p. Neuronal regions of interest were curated after visual inspection of their shape and fluorescence signals. Neuronal fluorescence signals were consolidated by upsampling the signals obtained with the lowest frame rate to match those acquired with the maximum frame rate. To relate neuronal activity to behavior, the consolidated fluorescence signals were upsampled again to match the framerate of the behavior video.

### Alignment of 2-photon images to Allen Common Coordinate Framework

Prior to 2-photon imaging experiments, the locations of the forelimb and hindlimb somatosensory cortex were determined by sensory mapping. Sensory mapping experiments were conducted under similar conditions as the single-photon imaging experiments, only mice were anesthetized with 1.2% isoflurane in oxygen and images were acquired at 40 fps (Figure 4-figure supplement 1D). Piezoelectric stimulators touching the forelimb or hindlimb were used to record somatosensory-evoked stimulus responses (Y. Xie et al. 2016). Averages of responses to sensory stimulation were calculated from 20 trials of stimulation with an interstimulus interval of 10 s (Figure 4-figure supplement 1E). After each 2-photon imaging session, a 5×5mm^2^ image of the cortical surface was taken with the Diesel2p’s linear scanners. ROIs captured with the resonant scanners were registered to this larger field-of-view 2-photon image, which was then registered to a 1024×1024 pixel template image (8.2×8.2mm^2^), obtained on the single-photon imaging system (Figure 4-figure supplement 1A-C). The single-photon template image was then registered to the dorsal projection of the Allen Common Coordinate Framework using control points placed in the mapped somatosensory cortices (Figure 4- figure supplement 1F). Each image registration process yields a transformation matrix, which was used to transform each neuron’s pixel location in the resonant image obtained from Suite2p, to its location in the Allen Common Coordinate Framework.

**Figure 4.**
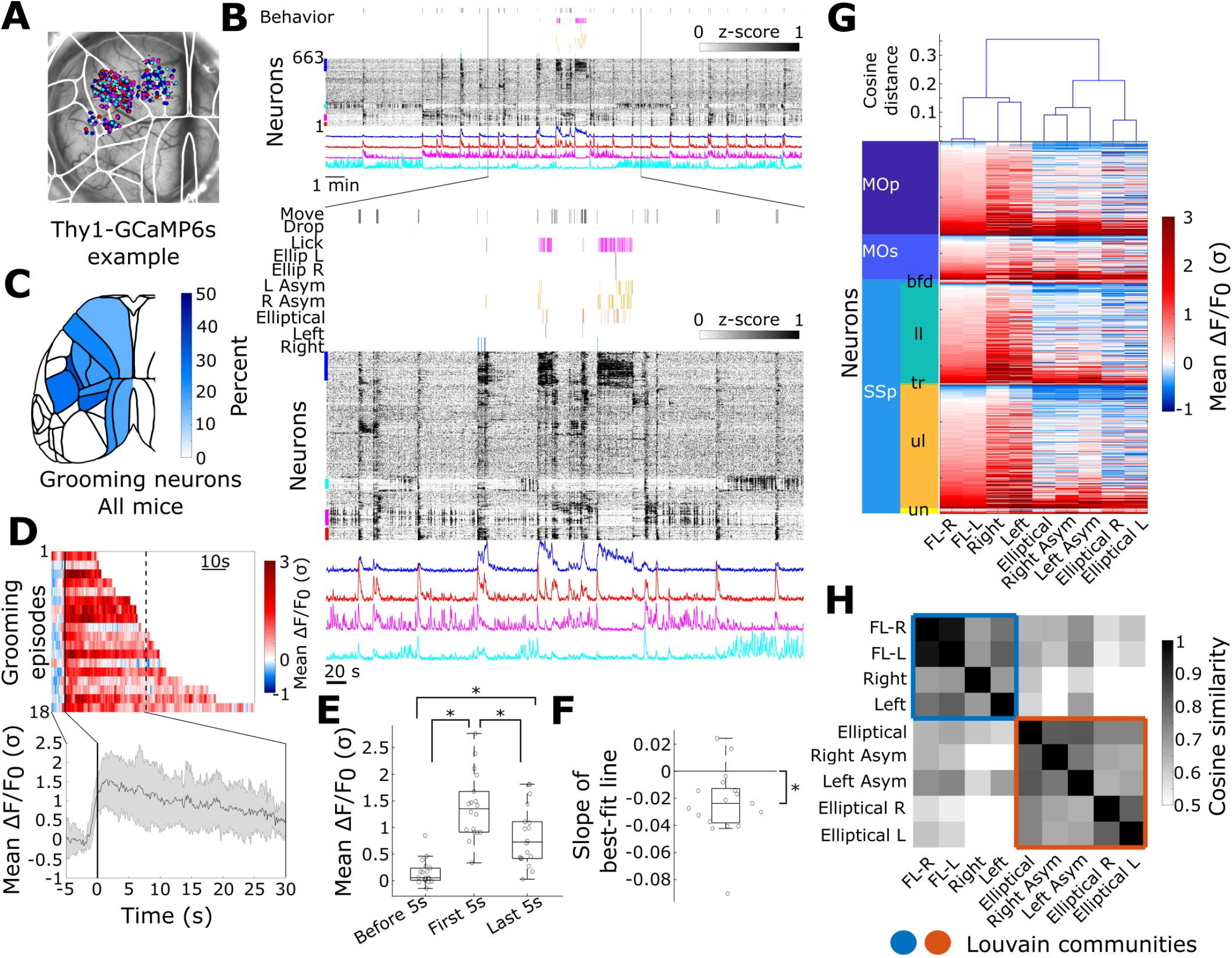
Grooming-responsive neurons observed across cortical areas with 2-photon imaging. **A)** Example session showing neuron locations in the cortical atlas. This session corresponds to examples shown in panel (B) and (G). Neurons are color coded by the populations highlighted in (B). Black neurons are unhighlighted population. **B)** Top: Rastermap embedding of neuronal fluorescence activity with behavior ethogram shown above. Colored lines on the left side of the rastermap show highlighted neuronal populations. Average of the highlighted populations are shown below. Bottom: Expanded view of the data. **C)** Percentage of imaged neurons that belong to the grooming-responsive population observed in each cortical region. **D)** Top: Heatmap showing averaged fluorescence across the grooming-responsive population of neurons, for each episode exceeding 10s in duration. Dotted line is drawn at 50th percentile of episode duration. Bottom: Averaged fluorescence across all grooming episodes. **E)** Comparison between averaged cortical signals at various time points. All time points significantly different. n=18 grooming episodes across 3 mice. p<0.05 repeated measures ANOVA with post-hoc Bonferroni correction. **F)** Slope of the best fit line for all grooming episodes exceeding 10s. Slope is less than 0. p<0.05 t-test. **G)** Heatmap of averaged neuronal fluorescence during each behavior. Cells are sorted by cortical region and responses to non-grooming forelimb movements. Data are from the example mouse shown in panels (A) and (B). Top: Dendrogram showing cosine distance between neuronal responses across behaviors. **H)** Averaged cosine similarity matrix across all mice. Louvain community groupings are denoted by colored squares.

### 2-photon image analysis

Neurons that exhibited overlapping activation with grooming episodes were extracted after visualizing the Rastermap embedding (Stringer et al. 2024) of neuronal activity with overlaid behavior ethograms (Figure 4B). The identified population of neurons was verified by computing the Pearson correlation coefficient between each neuron’s fluorescence activity with binarized grooming and non-grooming forelimb movement signals (Figure 4-figure supplement 2A-B). Binary grooming signals were generated by combining the individual grooming component behaviors and applying a 3-second aggregation window, as previously described. To extract non-grooming forelimb movement signals, paw velocities tracked with DeepLabCut were binarized by assigning a value of 1 to velocities exceeding the session-wide mean plus one standard deviation, and 0 otherwise. This produced a binary signal indicating periods of forelimb movement versus stillness. Grooming periods were then excluded from this signal, yielding a binary trace of non-grooming forelimb movements.

To assess differences in neuronal responses associated with different behaviors, neuron activity was averaged during all frames concurrent with the respective behaviors. The similarity between neuronal responses to behaviors were assessed using the cosine distance, which measures the angle between neuronal response vectors and therefore groups behaviors by the directionality of the responses across the neuronal population. All subsequent results were comparable using the correlation distance as well. To determine the similarity of neuronal responses to different behaviors across mice, the consensus similarity matrix was computed as the average cosine similarity matrix across all mice and sessions (n=9 sessions across 4 mice). In 3 out of 9 sessions, mice did not exhibit all grooming behaviors, resulting in a sparse similarity matrix containing NaN values, as distances could not be computed between behaviors which were not observed. The consensus similarity matrix was therefore computed as the average across all mice, ignoring NaN values found in the sparse similarity matrices. Louvain community detection (Rubinov and Sporns 2011) was then performed on this consensus matrix, as well as on each of the individual sessions in which all of the behaviors occurred (n=6 sessions across 3 mice), and the adjusted rand index was computed between individual sessions and the consensus session.

### Statistics

Data were first checked for normality using a Kolmogorov-Smirnov test. If all groups were normal, comparisons between two groups were assessed using a t-test, and comparisons between multiple groups were assessed with a repeated measures ANOVA with a post-hoc Bonferroni correction for multiple comparisons. If data were not normally distributed, comparisons between two groups were made with a Wilcoxon rank-sum test. Throughout the manuscript, all data are reported as mean +/- standard deviation with the exception for the data reported in Figure 1-figure supplement 2 and Figure 4-figure supplement 2A, which is reported as median +/- interquartile range; and Figure 3-figure supplement 3B, which shows the mean +/- standard error.

### Resource availability

Code for preprocessing and analysis are available at: https://github.com/ubcbraincircuits/innate-behaviors. Data are available on the Open Science Framework at https://osf.io/fa2ut/.

## Results

### Evoked grooming behaviors demonstrate rich diversity in movements

Behaviors were monitored with a camera placed directly in front of the mouse, facilitating tracking of the paws with DeepLabCut (Mathis et al. 2018) (Figure 1A). Grooming behaviors were stimulated by placing a water drop onto the whisker pad, once per minute over the course of 20 mins (Figure 1B). Spontaneous and stimulated grooming behaviors were assessed (Figure 1C). 44 paw movement trajectory features were extracted for each grooming behavior (Table 1). This feature matrix was projected into a 2-dimensional space using UMAP (Figure 1D). Hierarchical clustering on these embeddings separated the data into seven groups (Figure 1D, silhouette score = 0.42), which represent distinct behaviors as observed by the corresponding movement trajectories (Figure 1E and Figure 1- figure supplement 1C) as well as strong agreement with manually annotated behaviors (adjusted rand index = 0.6758; adjusted mutual information score = 0.6759, Figure 1-figure supplement 1A-B). The behaviors detected here largely agree with previously reported grooming component behaviors (Kalueff et al. 2016; Cromwell and Berridge 1996), however some differences are worth noting: 1) The elliptical asymmetric behaviors observed here have not been described elsewhere. 2) Due to the lateralized nature of the stimulus, pure unilateral grooming strokes, which are not part of the typical grooming repertoire, occur frequently in this paradigm. (Note that in prior naming conventions, unilateral grooming strokes involve both paws - these actions are referred to as Asymmetric in this work). 3) Due to the head-restrained nature of the experimental setup, grooming chains do not typically progress to large bilateral strokes or body licking actions, as this necessitates movement of the head. In rare cases, mice attempted large bilateral strokes (Figure 1figure supplement -1B-C), but these events were too infrequent to be resolved with the clustering methods employed here and could not be analyzed with sufficient statistical power.

Water drop evoked sessions resulted in a significant increase in the number of grooming episodes compared to spontaneous sessions where no drop was delivered (Figure 1F-G, p=2.1×10^−12^, t-test). While grooming behaviors tend to align with the delivery of the water drop stimulus, prolonged grooming episodes can be observed throughout the course of the session (Figure 1F). Additionally, evoked grooming sessions altered the composition of grooming behaviors, increasing the relative frequencies of unilateral and asymmetric bilateral grooming actions (Figure 1H). Most of these actions, with the exception of the Elliptical Asymmetric behaviors (Figure 1E), were significantly dependent on the placement of the water drop stimulus (Figure 1-figure supplement 2. For right sided stimuli - right vs left unilateral 21+/-28 vs 3+/-4, p=8.1×10^−6^ Wilcoxon rank-sum test; right asymmetric vs left asymmetric 10+/-27 vs 3+/-23, p=0.0081, Wilcoxon rank-sum test; elliptical right vs elliptical left 4+/-7 vs 3+/-11, p=0.37 t-test. For left sided stimuli - right vs left unilateral 3+/-4 vs 23+/-11, p=0.039 t-test; right asymmetric vs left asymmetric 30+/-20 vs 52+/-25, p=0.17 t-test; elliptical right vs elliptical left 19+/-9 vs 13+/-12 p=0.18 t-test. All data reported as median +/- interquartile range).

### Head-restrained grooming episodes are variable, but organized

Natural grooming is a serially ordered behavior, which often presents in a stereotyped rostrocaudal progression beginning with grooming actions directed towards the nose (elliptical strokes), progressing to larger asymmetric actions directed towards the face (asymmetric strokes), then bilateral strokes which over the ears, and finally terminating with licking directed towards the body. This progression is termed the syntactic chain (Kalueff et al. 2016; Cromwell and Berridge 1996). While the head-restrained nature of the experimental setup limits the progression of the syntactic chain to the first 2 phases (i.e. Elliptical and Asymmetric strokes), atypical grooming behaviors such as unilateral and elliptical asymmetric strokes were observed (Figure 1H). To determine if the head-restrained grooming behaviors exhibit a similar phasic organization as observed in naturalistic grooming, their temporal presentation within grooming episodes were assessed. Grooming episodes varied in their duration and composition (Figure 2A-B, episode duration range: 0.12-110.16 seconds). Short duration grooming episodes were the most common (Figure 2B), which mostly consisted of unilateral grooming actions. However, during longer grooming episodes, unilateral grooming behaviors make up a small proportion of the total number of actions performed (Figure 2C). Analysis of the transitions between component behaviors within episodes revealed closer associations between certain grooming behaviors as determined with Louvain community detection (Figure 2D-E). Transition matrices contained states which represent each of the classes of grooming behaviors identified in Figure 1, as well as Start and Stop states, which represent the period immediately preceding or following a grooming episode, respectively. Licking behaviors were excluded from transition analyses, as these behaviors often occur simultaneously with the other grooming actions (Figure 2- figure supplement 1A-B). Louvain community assignments were robust to parameter selection (e.g. aggregation window size and gamma parameter in Louvain community detection algorithm) and consistent across individual mice (n=8 mice; adjusted rand index = 0.93 +/- 0.066 (mean +/- standard deviation)). Unilateral grooming actions reliably clustered with Start and Stop states (Figure 2D-E), suggesting that unilateral actions are often stand-alone events that are not typically embedded within more complex grooming behavior chains (Figure 2C-E). Note that the majority of sessions were conducted with the stimulus directed toward the right side, yielding fewer left unilateral actions (Figure 1-figure supplement 2). On the other hand, elliptical, asymmetric strokes and elliptical asymmetric strokes reliably cluster separately, and have little association with Start and Stop states (Figure 2D), suggesting that these actions reflect the distinct phases in grooming behaviors that have been previously reported.

### Transient activation of cortical networks during continuous behaviors

Cortex-wide neuronal activity during grooming behaviors were assessed with mesoscale single photon calcium imaging using GCAMP expressing mice (Figure 3A). Large cortical calcium events were associated with the onset of grooming behaviors (Figure 3B), but sustained grooming episodes exhibited a gradual decline in cortical activation throughout the behavior (Figure 3C-F). Averaged cortical activation during grooming episodes exceeding 10 seconds in duration were significantly greater at the onset of the behavior compared to late in the episode or before the episode began (Figure 3E, n=56 grooming episodes, repeated measures ANOVA, F(2,108)=41.8, p=3.5×10^−14^, with post-hoc Bonferroni correction; onset vs late: 0.86+/-0.64 vs 0.08+/-0.44, p=9.17×10^−11^; onset vs before: 0.86+/-0.64 vs 0.04+/-0.48, p=7.5×10^−9^; before vs late: 0.04+/-0.48 vs 0.08+/-0.44, p=1). Additionally, the slope of the line which best fit the averaged cortical activity throughout the duration of the grooming episode was significantly below zero indicating a progressive decrement (Figure 3F, average slope: -0.058+/-0.066, one-sample t-test, p=1.24×10^−8^, t=-6.5). Individual behaviors were associated with activation of distinct cortical networks (Figure 3G). For example, licking actions were associated with activation of anterior-lateral portions of the motor cortex (Figure 3G), while unilateral actions were associated with activation of contralateral cortical networks spanning whisker, forelimb, and hindlimb somatosensory networks, as well as the medial portion of secondary motor cortex (Figure 3G). Right and left asymmetric grooming actions were similarly associated with contralateral cortical network activation, although on average, frontal cortical activation was less pronounced (Figure 3G). Elliptical and elliptical asymmetric grooming behaviors exhibited bilateral activation patterns. In general, the maximally activated networks involved in licking and unilateral grooming behaviors appeared to be the most consistent across animals compared to the bilateral grooming movements (Figure 3G). Averaged cortical activation maps associated with licking and elliptical behaviors were qualitatively similar between evoked and spontaneous sessions, where the water drop was not applied (Figure 3-figure supplement 1). However, most grooming behaviors were too infrequently expressed in the spontaneous context to draw meaningful comparisons (Figure 3-figure supplement 1, Figure 1F-H).

Because grooming behaviors, particularly the bilateral actions, often occur in quick succession with each other and simultaneously with licking behaviors (Figure 2A), a linear ridge regression model (Musall et al. 2019) was employed to unmix cortical responses associated with each behavior (Figure 3- figure supplement 2A-D). All grooming behaviors were included in the model (Figure 3-figure supplement 2A), as well as the timing of the water drop stimulus, the audio cue, and non-grooming movements. These binary regressors were used to create a design matrix, which was fit to the neural fluorescence data using ridge regression with 10-fold cross validation (Musall et al. 2019), yielding time-varying coefficient weight maps for each behavior (Figure 3-figure supplement 2B). Behavior kernels appeared qualitatively similar to the averaged responses (Figure 3G, Figure 3-figure supplement 1-2B), and the variance explained across the cortex was greatest along the forelimb and hindlimb somatomotor cortex (Figure 3-figure supplement 2C). To evaluate each behavior’s unique contribution to the model, a reduced model was created for each behavior by randomly permuting the behavior of interest, and recomputing the regression. The unique contribution was then calculated as the difference in explained variance between the full model and the reduced model (Figure 3-figure supplement 2D). Interestingly, although the maximally activated networks during licking behaviors were associated with anterior lateral cortical areas (Figure 3G, Figure 3-figure supplement 2D), the greatest contribution to explained variance from licking kernels consistently involved posterior networks including trunk and hindlimb somatosensory cortex and posterior parietal areas (Figure 3-figure supplement 2D). This was associated with the large negative coefficients in the posterior cortex associated with licking (Figure 3- figure supplement 2C). The unique contribution from unilateral behaviors matched the networks determined by the averaged maps (Figure 3-figure supplement 2D, Figure 3G). Similarly, the asymmetric grooming actions involved contralateral sensory networks along the lateral edges of the cortex (Figure 3-figure supplement 2D). Generally, the most consistent networks identified using the ridge regression approach appeared to be those related to unilateral grooming actions and licking behaviors, similarly to those identified by the averaged activation maps (Figure 3-figure supplement 2D, Figure 3G).

To determine if the transient cortical activation observed at the onset of prolonged grooming bouts is unique to grooming movements, and to clarify the contribution of licking behaviors to mesoscale cortical network activity, widefield cortical calcium activity was imaged while thirsty mice drank water over extended periods of time (Figure 3-figure supplement 3A-B). Licking behavior was continuously expressed over the period when water was delivered (Figure 3-figure supplement 3B). Averaged cortical activity maps during licking showed activation of the anterior cortex with similar networks activated as in the grooming context (Figure 3-figure supplement 3C, Figure 3G). Additionally, transient cortical activation was observed at the onset of continuous drinking but subsided over time (Figure 3-figure supplement 3D). Movements of the paws, however, were associated with large cortical calcium events even when expressed throughout the drinking period (Figure 3-figure supplement 3D). Cortical responses associated with each behavior were evaluated with ridge regression as described earlier. In agreement with the averaged maps and expected networks, time-varying coefficient maps for licking and paw movements highlight responses in the anterior motor cortex and hindlimb somatosensory cortex, respectively (Figure 3-figure supplement 3E-F). The unique explained variance maps for the licking behavior differed in the drinking context compared to the grooming context (Figure 3-figure supplement 3F).

### Single-cell activity

Single layer 2/3 excitatory neuron responses to grooming behaviors were assessed across cortical areas using the Diesel2p wide field-of-view 2-photon microscope (Figure 4A). The anatomical location of all neurons was estimated by registering imaging data to the Allen Institute Common Coordinate Framework (Figure 4-figure supplement 1). Cortical neurons exhibited rich dynamics throughout the session, as shown by the Rastermap embedding in an example mouse (Figure 4B). In the example session (Figure 4B), neuronal populations that were active during grooming (blue), active during movements but not grooming (red), basally active but suppressed during grooming (magenta), and anticorrelated with the penultimate population (cyan) were observed (Figure 4B). These neuronal populations were distributed across cortical regions (Figure 4A). All sessions in which extended bouts of grooming behavior occurred (n=7 sessions over n=4 mice) exhibited populations of neurons which were responsive to grooming behaviors, as identified with Rastermap. This neuronal population exhibited significantly higher Pearson correlation coefficients with grooming behaviors compared to the remaining neuronal population (Figure 4-figure supplement 2A, grooming neurons (n=418) vs all other neurons (n=2198): 0.33+/-0.24 vs 0.030+/-0.087 [median +/- interquartile range]; Wilcoxon rank-sum test, p=4.5*10^−198^), and correlation coefficients with grooming behaviors were significantly higher within this neuronal population compared to those with non-grooming movements (Figure 4-figure supplement 2B, grooming vs non-grooming movements: 0.34+/-0.16 vs 0.089+/-0.067; paired t-test, p=8.8×10^−107^, t=30.1). These grooming-responsive neurons were distributed across all cortical areas imaged (Figure 4C). To determine if fluorescence signals decline over long periods of grooming as was observed in the single-photon data, the averaged fluorescence across the grooming-responsive population of neurons were assessed for all grooming episodes with a duration exceeding 10s (Figure 4D). The averaged activity at the end of the grooming episode was significantly lower than at the beginning but still greater than the baseline period before the grooming episode starts (Figure 4E, n=18 grooming episodess; repeated measures ANOVA F(2,36)=47, p=1.0×10^−10^, with post-hoc Bonferroni correction; first 5s vs last 5s: 1.5+/-0.72 vs 0.92+/-0.64, p=8.8×10^−5^; last 5s vs before 5s: 0.92+/-0.64 vs 0.19+/-0.27, p=1.9×10^−4^; before 5s vs first 5s: 0.19+/-0.27 vs 1.5+/-0.72; p=3.9×10^−7^). Similarly, the slope of the best-fit line was also significantly below zero (Figure 4F, average slope: -0.024+/-0.025 one-sample t-test: p=2.9×10^−4^, t=-4.2). While a sizable population of neurons had activity correlated with grooming (Figure 4C), individual grooming behaviors did not have distinct neuronal representations, with the exception of unilateral grooming actions, which more closely resembled non-grooming forelimb movements (Figure 4G). Louvain community detection (Rubinov and Sporns 2011) on the averaged similarity matrix across all sessions and mice (n=9 sessions, n=4 mice) yielded two groups separating unilateral grooming behaviors and non-grooming forelimb movements from all other bilateral grooming behaviors (Figure 4H). This grouping was consistent across individual mice and sessions where all behaviors were exhibited and the similarity matrix could be generated (n=6 sessions, n=3 mice, adjusted rand index = 0.87+/- 0.13 [mean +/- standard deviation]).

## Discussion

Grooming behaviors are ubiquitous across animals (Fentress 1968b, 1968a; Kalueff et al. 2016) serve a variety of physiological functions such as removal of dirt or parasites, thermoregulation, and social communication (Feusner, Hembacher, and Phillips 2009). Excessive and pathological self-grooming is observed in humans with obsessive-compulsive or related disorders such as body dysmorphic disorder (American Psychiatric Association 2021) autism spectrum disorder (Peça et al. 2011) obsessive compulsive disorder (Welch et al. 2007). Assessing neural activity during grooming behaviors in rodents may therefore provide translational value in understanding the neural circuit mechanisms involved in pathological grooming behaviors associated with human psychiatric disorders (Kalueff et al. 2016; Burguière et al. 2013). Moreover, because grooming behaviors in rodents are complex, highly stereotyped, and sequentially patterned motor actions, grooming presents a model for studying the neural mechanisms of motor control underlying such behaviors (Kalueff et al. 2016)(Tartaglione et al. 2016).

We assessed grooming behavior in head-restrained mice, and the cortical responses associated with these behaviors at mesoscale and layer 2/3 single-cell resolution. Head-fixed grooming behaviors were reliably evoked using a water drop stimulus on the whisker pad. These behaviors include stereotyped bilateral grooming actions, which were temporally organized in such a way that transitions within classes of behaviors (i.e. [1] elliptical; [2] right or left asymmetric; and [3] elliptical right or elliptical left) were more likely to occur than transitions between classes. The separate grouping of these behavioral classes by Louvain clustering resembles the distinct phases observed in stereotyped spontaneous grooming (Cromwell and Berridge 1996), suggesting that the patterned structure of grooming behaviors is preserved in the head-restrained context, despite the fact that head-restraint limits the behavioral repertoire to actions directed towards the face and nose. The head-restrained grooming paradigm could therefore be used with multimodal methods to study cortico-striatal interactions (Ramandi et al. 2023; Peters et al. 2021) during striatal patterning of innate sequentially ordered behaviors.

In addition to the stereotyped, bilateral paw movements, unilateral strokes were frequently observed, which were directed towards the side of the stimulus and do not typically occur during spontaneous grooming sessions. These unilateral strokes also clustered separately from the other grooming behaviors based on their temporal presentation, typically occurring either independently or before a longer stereotyped grooming episode. Unilateral grooming actions were associated with consistent mesoscale cortical activation spanning contralateral sensory and motor networks, and at the cellular level, were more similar to non-grooming forelimb movements than to bilateral grooming behaviors. One possible explanation for these disparities is that the movements during unilateral grooming actions reflect voluntary forelimb movements, while the bilateral grooming actions reflect automatic, subcortically-driven behaviors. In line with this, parallels have been drawn between unilateral grooming actions and reaching for food (Naghizadeh, Mohajerani, and Whishaw 2020), which is a cortex-dependent action (Galiñanes, Bonardi, and Huber 2018; Yang, Kanodia, and Arber 2023) that engages similar cortical networks (Y. Wang et al. 2023). Further experiments employing inhibition of contralateral sensory and motor cortical networks during this evoked grooming paradigm are necessary to determine if the unilateral grooming movements observed here are cortex-dependent behaviors.

While stereotyped bilateral grooming behaviors are not cortex-dependent (Ruder et al. 2021; Guo et al. 2015; Berridge 1989), grooming-responsive neurons were observed across all mice and in all cortical regions imaged. This population of neurons exhibited activation that persisted throughout the duration of prolonged grooming episodes regardless of the class of behavior, and was significantly more correlated to grooming behaviors than non-grooming forelimb movements. At the mesoscale level, spatial patterns of cortical network activation during bilateral grooming behaviors did not appear as consistent as during unilateral behaviors. Nevertheless, activation was observed in the posterolateral sensory networks spanning whisker, nose, and visual cortex. The activation of these regions may simply reflect passive sensory responses associated with the stimulus or grooming behaviors (e.g. sensation of the drop on the vibrissae, or the paw brushing against the whisker pad). Alternatively, since brainstem regions responsible for grooming receive input from somatosensory cortex (Z. Xie et al. 2022), lateralized activation of sensory cortex may shape grooming behaviors, for example, to incorporate more asymmetric behaviors directed towards the contralateral side where the stimulus was delivered. Further experiments using lateralized optogenetic stimulation during grooming would be required to determine if cortical network-level activation has a functional impact on the expression of the behavior.

Prolonged grooming episodes were accompanied by an initial increase in cortical activity which reduced over time despite continuous expression of the behavior. Additionally, global cortical activity declined over extended periods of continuous drinking, and a decline in cortical activity has also been reported during prolonged bouts of locomotion (Vinck et al. 2015; West, Gerhart, and Ebner 2024). Grooming, drinking, and locomotion are all behaviors that can be expressed in decerebrate rodents, indicating that they are mediated by subcortical systems (Berntson and Micco 1976). The declining cortical responses may therefore suggest that mesoscale cortical activity mostly reflects the initiation of subcortically-mediated behaviors, rather than the behavior itself. Nevertheless, at the single-cell level, a grooming-responsive population of neurons was observed. This neuronal population also demonstrated a gradual reduction in signal, which differed from other movement-responsive neurons that exhibited a transient response at the onset of grooming, but remained inactive throughout the episode. The more severe decline in signal over time observed with single-photon widefield imaging might therefore be attributed to the fact that the widefield imaging integrates signal from each of these populations of neurons. Alternatively, this may be attributed to differences in the imaging modalities, as widefield imaging may primarily reflect dendritic calcium activity, while the two-photon signals reflect somatic calcium activity. It is unclear to what degree the persistent activation of the grooming-responsive neuronal population might affect other cortical operations, but recent work has shown that neuronal responses to auditory stimuli are diminished during grooming (Morandell et al. 2024). Future work should investigate this more explicitly by evaluating multiple sensory modalities as a function of latency to both spontaneous and evoked grooming episodes.

We also examined cortical activation associated with licking behavior across grooming and drinking contexts. Consistent with previous findings (Komiyama et al. 2010), anterior and lateral cortical regions were activated during licking. Ridge regression modeling revealed strong negative components in somatosensory and posterior cortical areas during grooming, but not during drinking. This difference may be due to the gradual decline in somatosensory activity observed during prolonged grooming. Because grooming involves paw movements that engage the somatosensory cortex, the overlap with licking may cause the regression model to attribute the decreasing activity over time to the licking regressor, resulting in negative weights. In contrast, since paw movements are less common during drinking, somatosensory cortical activity remains more stable, and the regression model does not produce negative weights in these regions. These findings highlight limitations in using a linear model to dissociate cortical responses associated with repetitive, overlapping behaviors across prolonged timescales.

Limitations of this work include a relatively low number of animals for each of the single photon (n=4) and 2-photon (n=4) imaging studies, and an absence of hemodynamic signal correction in the single-photon study. Hemodynamic signals may contribute to the gradual decline in fluorescence over extended periods of behavior observed in the single-photon data, but they are unlikely to be a major driver of this decline, as the magnitude of the hemodynamic signal is much smaller than that of the fluorescence signals (Xiao et al. 2017; Michelson, Vanni, and Murphy 2019). Moreover, a gradual decline in signal was observed in the grooming-responsive population of neurons in the 2-photon imaging study, where hemodynamic artifacts are less of a concern. The observed decline in cortical activity over extended periods of grooming, along with the distinct cortical representations of unilateral and bilateral grooming behaviors, was consistent across imaging modalities. This consistency provides some confidence in the findings, despite the relatively small sample sizes.

In conclusion, cortical responses to evoked grooming behaviors were studied in a head-restrained context. Grooming behaviors included atypical, reach-like forelimb movements directed towards the site of the stimulus, as well as stereotyped component forelimb movements, which exhibited a phasic organization. Neuron activity across dorsal cortical areas was found to reflect stereotyped grooming behaviors, and at the mesoscale network level, cortical activation patterns were primarily localized to posterolateral areas. The functional consequences of these observed cortical responses on grooming behaviors or other cortical operations remains an open question. Additionally, averaged activity across the dorsal cortex was found to subside over the course of extended periods of grooming and drinking behaviors. The neurophysiological mechanism underlying this decline, and whether this phenomenon occurs during other instances of continuous movements over long durations, warrants further investigation.

## Acknowledgements

This work was supported by resources made available through the Dynamic Brain Circuits Research Excellence Cluster and the NeuroImaging and NeuroComputation (NINC) Core at the UBC Djavad Mowafaghian Centre for Brain Health (RRID:SCR_019086). Timothy H Murphy (THM) was supported by Canadian Institutes of Health Research (CIHR) Foundation Grants FDN-143209 and PJT-180631, and the Natural Sciences and Engineering Research Council of Canada (NSERC) Grant GPIN-2022-03723.

---

**Figure 1 – figure supplement 1:**
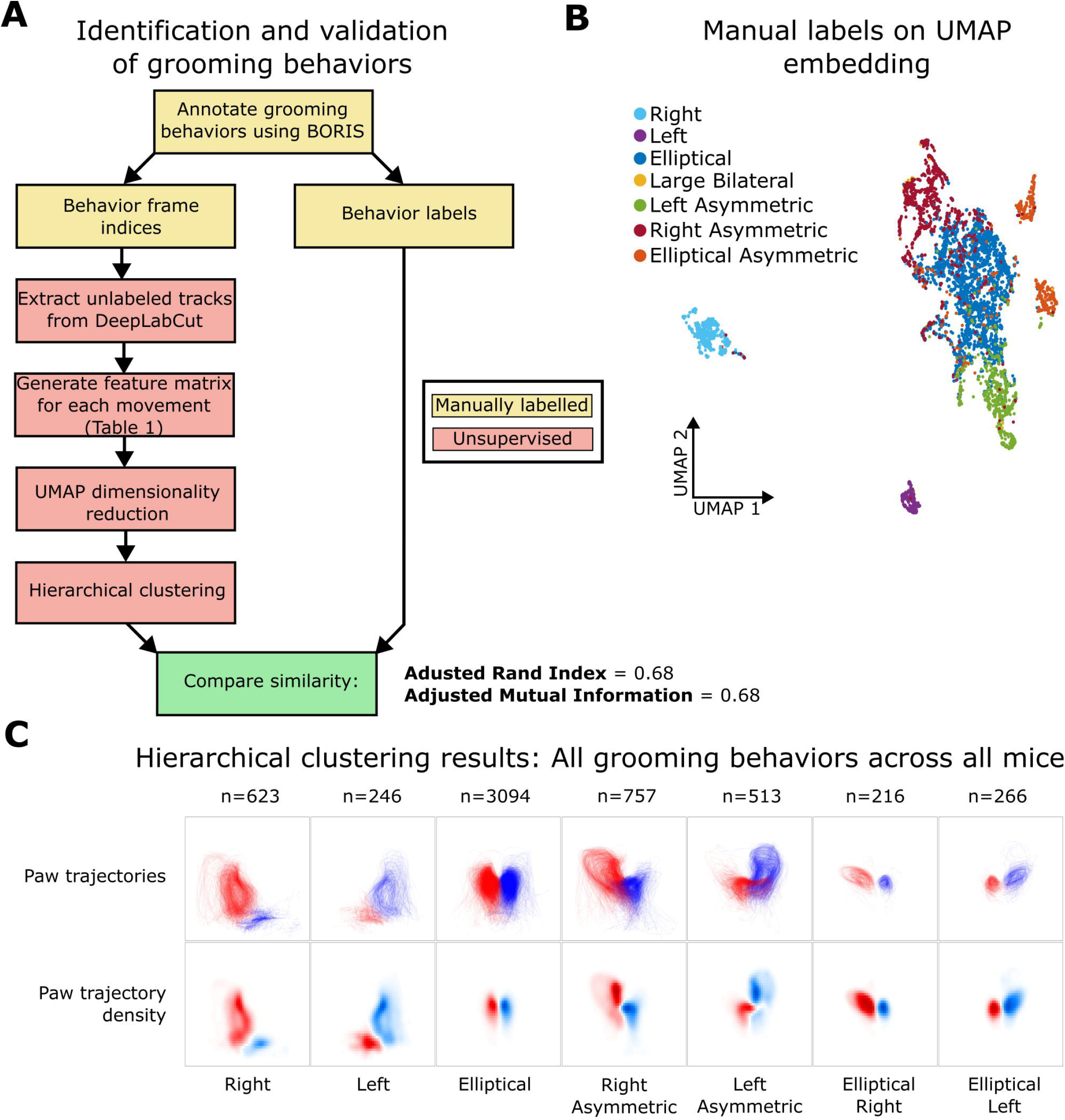
Methods for detecting grooming behaviors. **A)** Workflow for identifying and validating grooming behaviors. In the unsupervised method, features were generated from DeepLabCut tracks from keypoints on the paws (Figure 1A). **B)** Manual labels plotted in the same UMAP space as shown in Figure 1D. In the manual labelling scheme, large bilateral grooming events were classified as a separate group. These behaviors were extremely rare and were not categorized by the unsupervised method described in (A). **C)** Paw trajectories (top) and position probability density (bottom) for the right (red) and left (blue) paw, for all behaviors across all mice. The number of behaviors are displayed above.

**Figure 1 – figure supplement 2:**
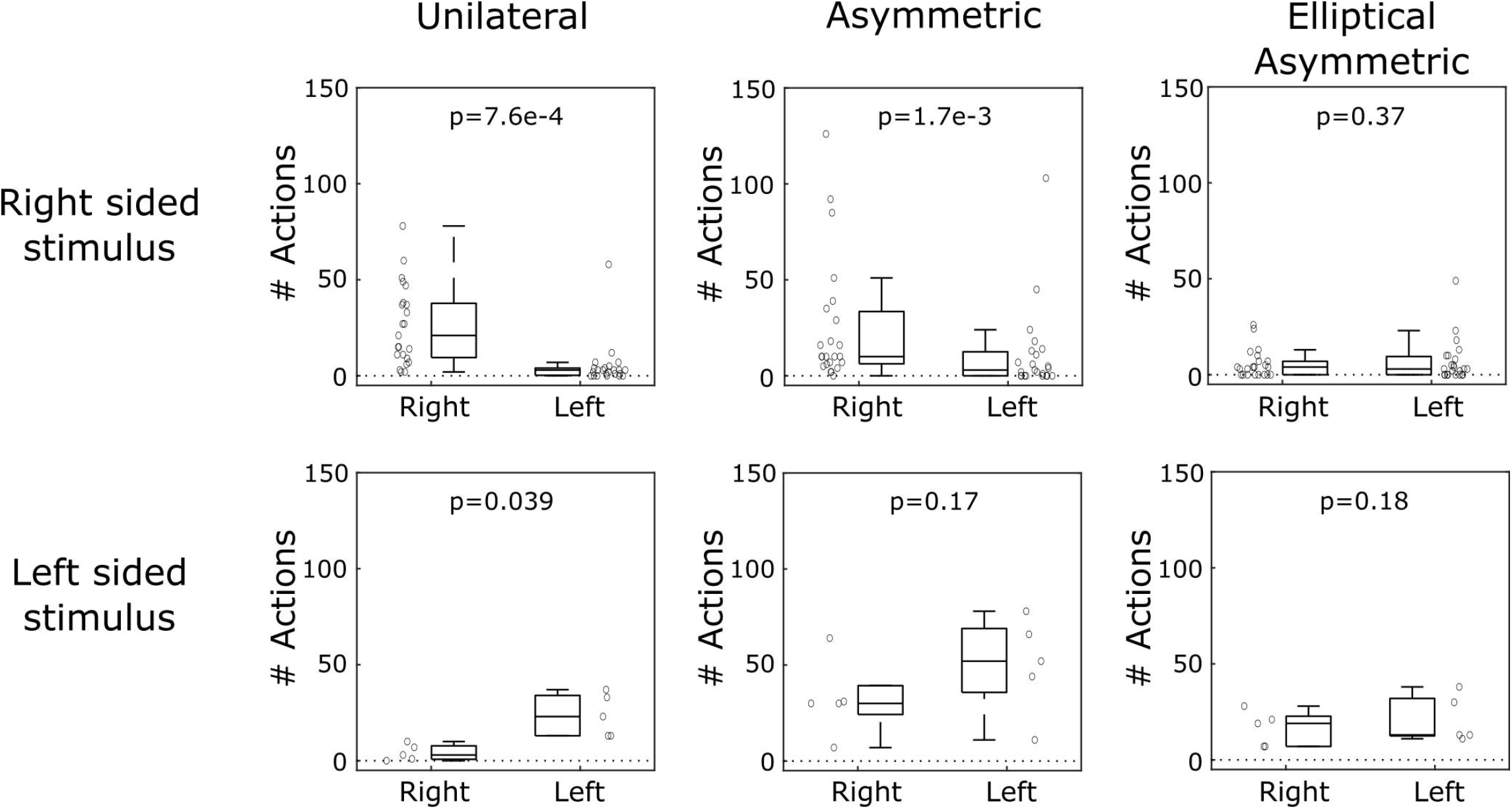
Lateralized water drop stimulus elicits lateralized responses. Drops directed towards the right side of the mouse significantly increased unilateral and asymmetric actions on the right side, but not elliptical asymmetric actions. Drops directed on the left side significantly increased unilateral actions on the left side, but did not result in significantly increased asymmetric or elliptical asymmetric actions directed towards the left.

Figure 1-Figure supplement 3: List of features from DeepLabCut tracks for grooming behavior identification

**Figure 2 – figure supplement 1:**
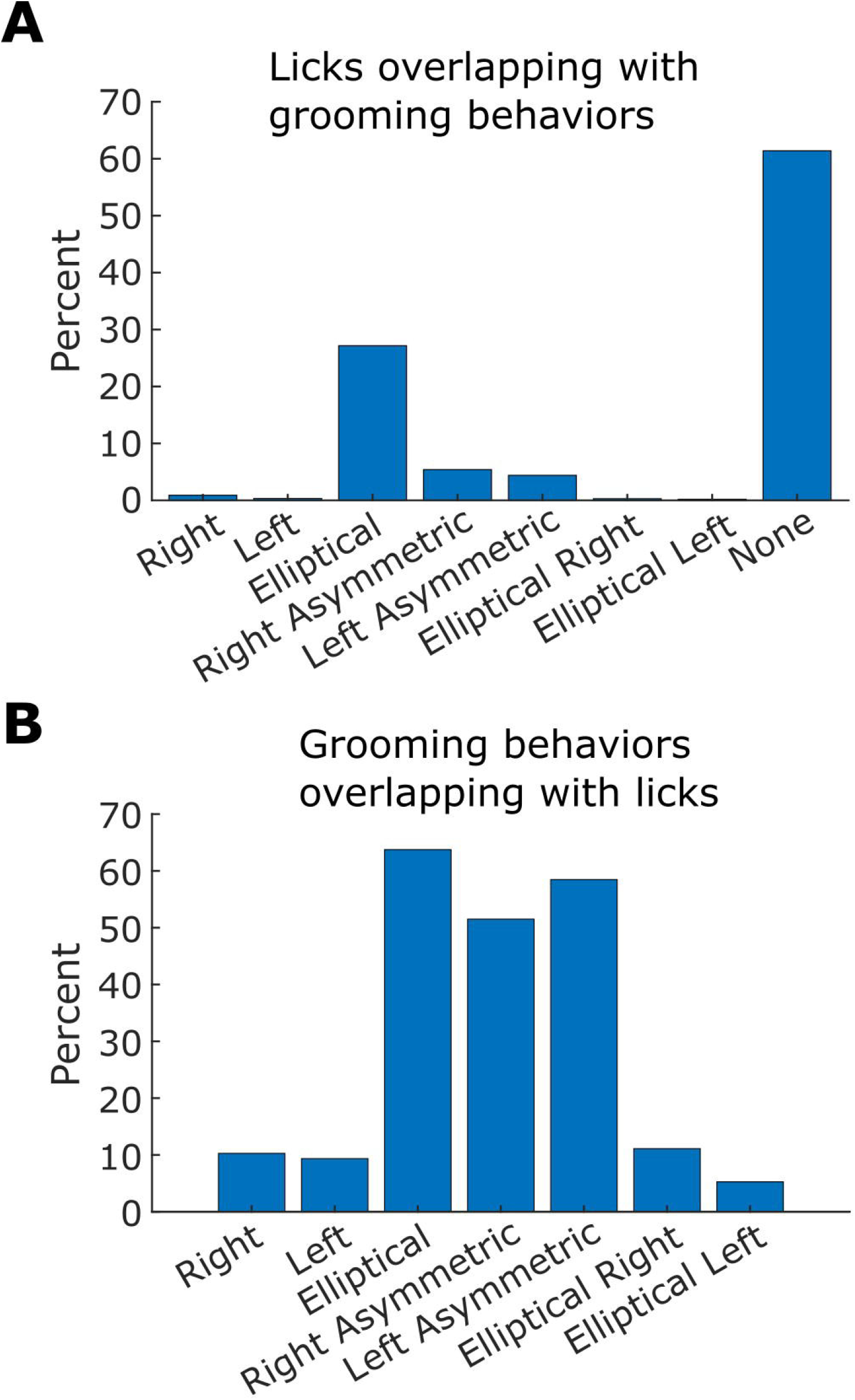
Licking and grooming behaviors are temporally overlapping. **A)** Percentage of lick events that overlap with each grooming behavior. **B)** Percentage of grooming behaviors that overlap with lick events. Data are compiled across all mice and sessions (n=7877 licks). N for individual grooming behaviors can be found in Figure 1 - figure supplement 1C.

**Figure 3 – figure supplement 1:**
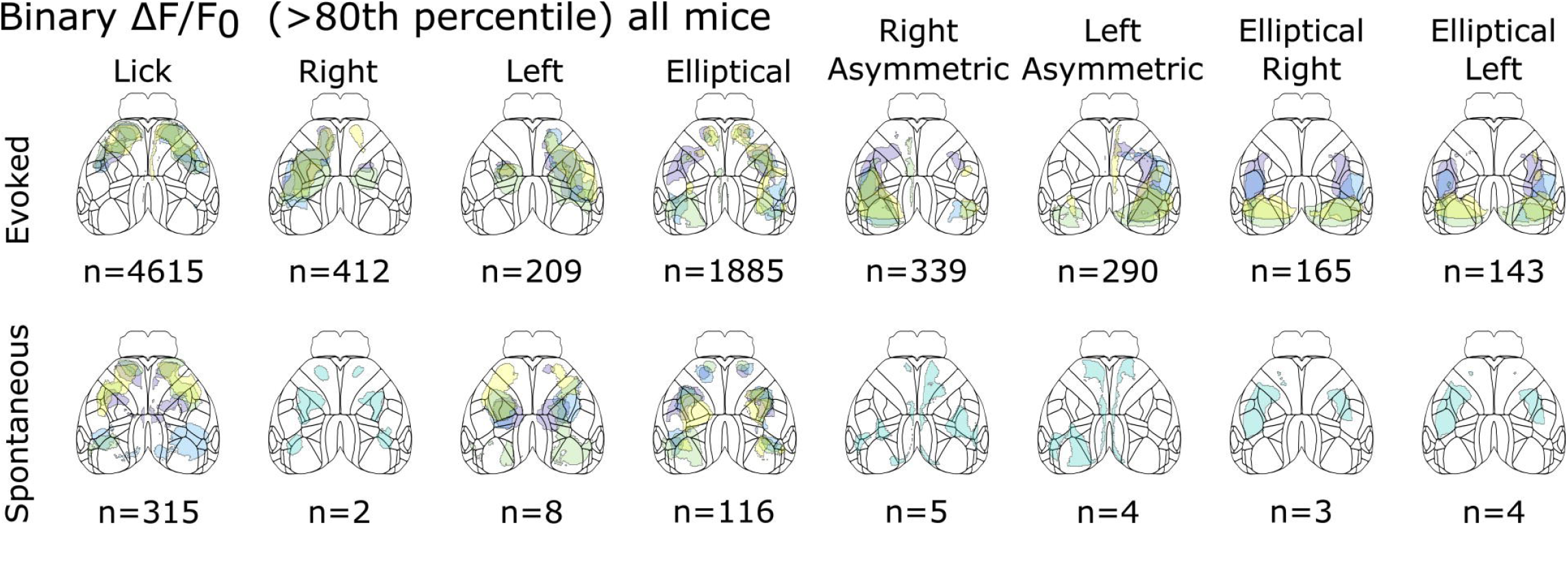
Cortical activation maps during spontaneous and evoked grooming behaviors. Binary response maps for each behavior are shown in the same format as in Figure 3G. Averaged response maps for each behavior are binarized at >80th percentile to show the maximally activated networks during evoked (top) and spontaneous (bottom) grooming behaviors. The total number of events across all animals are indicated below. Maps with only one colored contour indicate that only one mouse exhibited this behavior.

**Figure 3 – figure supplement 2:**
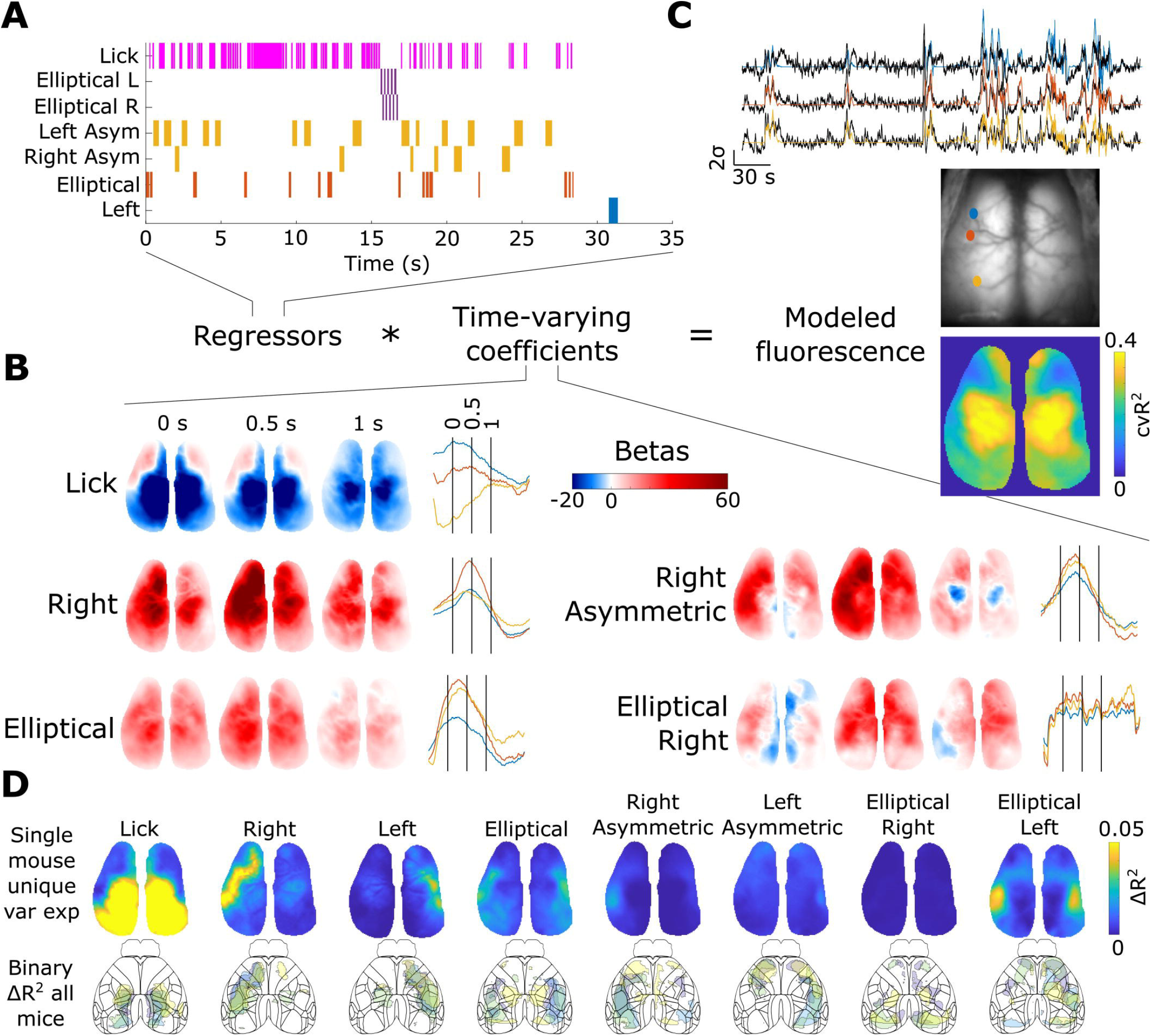
Regression analysis finds similar network involvement. **A)** Example ethogram of a single grooming event. The entire binary vectors for all behaviors are used as regressors in the ridge regression model. **B)** Time-varying coefficients solved for by the regression model. Each map shows the weights of the kernel at various time points. Time series data on the right show the weights as a function of time for cortical regions corresponding to those shown in (C). Vertical lines correspond to time points at which images are shown. **C)** Top: Example dFF traces from cortical areas (black) and model estimation (color). Middle: Cortical regions from where example dFF traces are shown in top. Bottom: Model explained variance across the cortical window over all sessions. **D)** Top: Unique contribution of each model variable to the total explained variance for an example mouse. Bottom: Unique contributions for each mouse and behavior binarized at >80th percentile to show maximally contributing cortical networks (bottom).

**Figure 3 – figure supplement 3:**
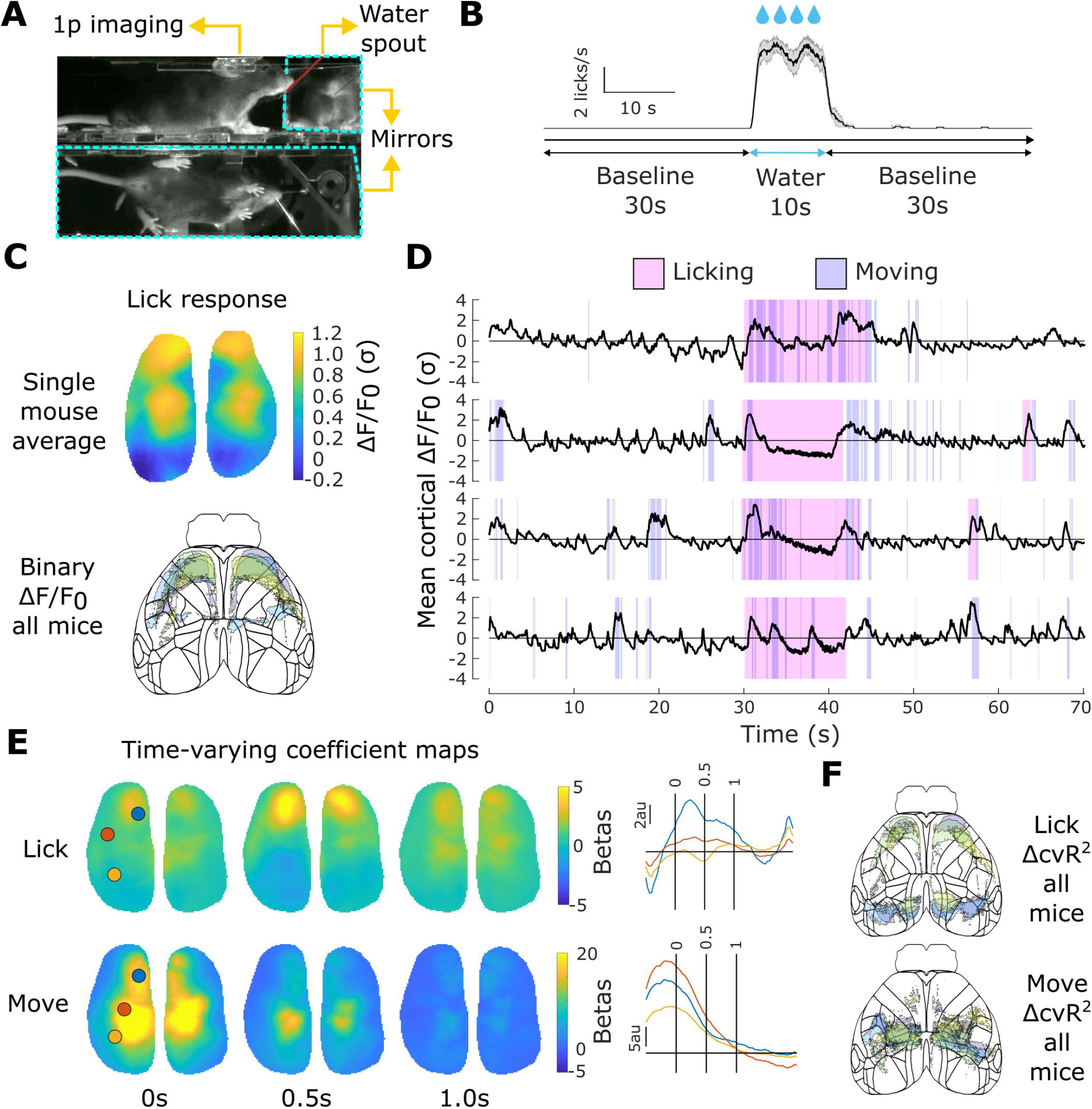
Transient mesoscale cortical network activation observed during continuous drinking behavior. **A)** Experimental setup. Mesoscale cortical calcium activity was imaged in water-restricted mice during continuous bouts of drinking. Movements of the paws and body were assessed with DeepLabCut. **B)** Trial structure. Following a brief baseline period, mice were given water for an extended period of time. Example licking rates from a single mouse over 12 sessions. Line and shaded areas indicate mean and standard error of the mean. **C)** Cortical activation during licking behavior in an example tTA-GCaMP6s mouse (top) and the maximally activated network across all mice (bottom, n=4). **D)** Averaged activity across the dorsal cortex from a tTA-GCaMP6s mouse over 4 example sessions. Licking and movement behaviors are annotated with magenta and blue shading, respectively. **E)** Time-varying coefficients solved for by ridge regression model for licking (top) and movement behaviors (bottom) in an example mouse. Each map shows the weights of the kernel at various time points. Time series data on the right show the weights as a function of time for the indicated cortical areas. **F)** Unique explained contribution for licking and movement variables to the total explained variance. Each mouse’s unique contribtions were binarized at >80th percentile to show the maximally contributing cortical networks.

**Figure 4 – figure supplement 1:**
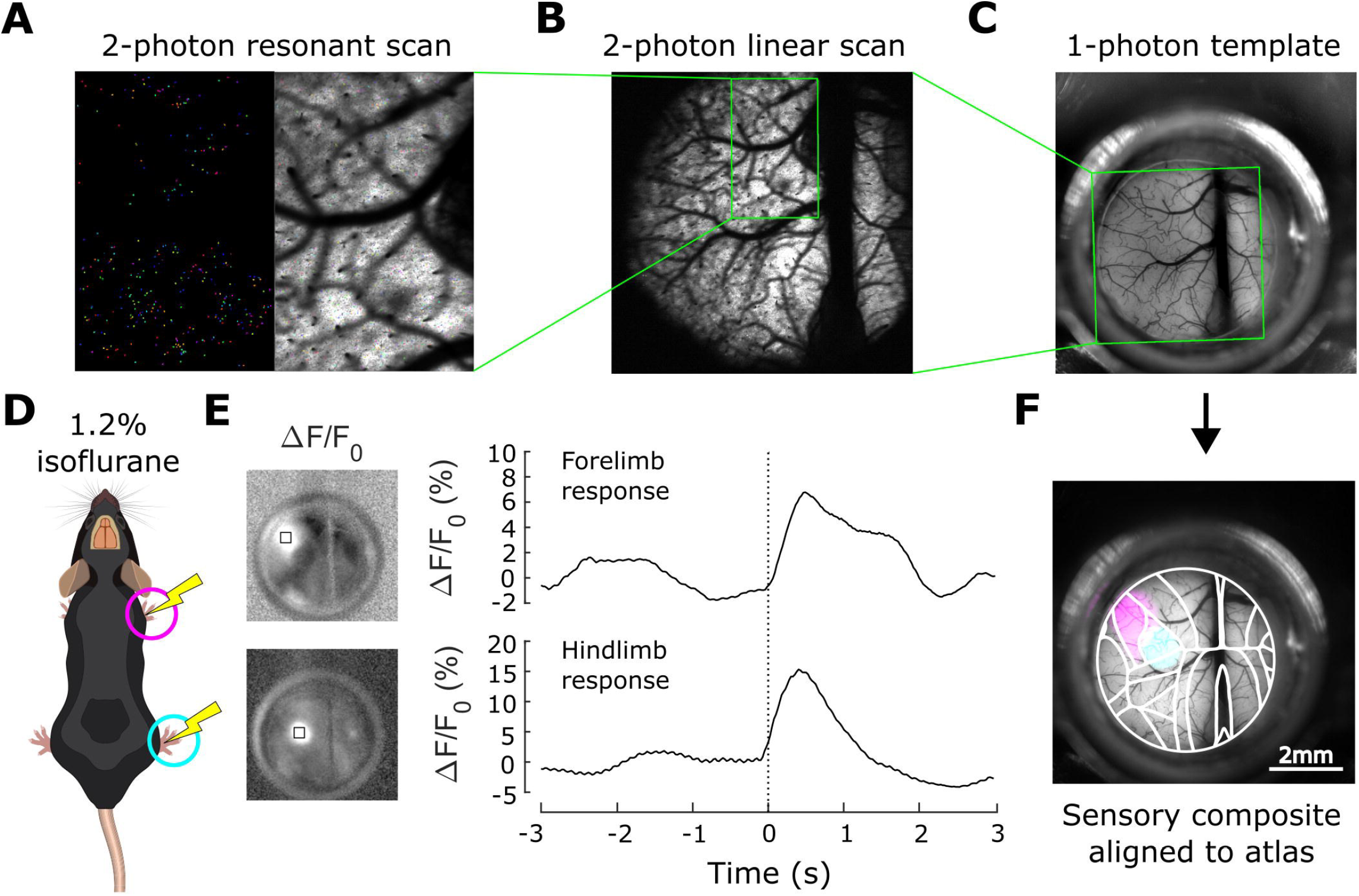
Alignment of 2-photon images to the Allen Institute cortical atlas. **A)** Example 2-photon resonant scan image showing neuron regions of interest detected with Suite2p. **B)** Example 2-photon linear scan image with larger field of view. Image from (A) was registered to (B). **C)** High resolution single-photon template image. Image from (B) was registered to (C). **D)** Protocol for mapping somatosensory cortex. **E)** Example response maps (left) and time courses within the indicated region from forelimb and hindlimb stimulation. **F)** Sensory response maps were overlaid on the high resolution template image, and used to register the template map to Allen Institute cortical atlas.

**Figure 4 – figure supplement 2:**
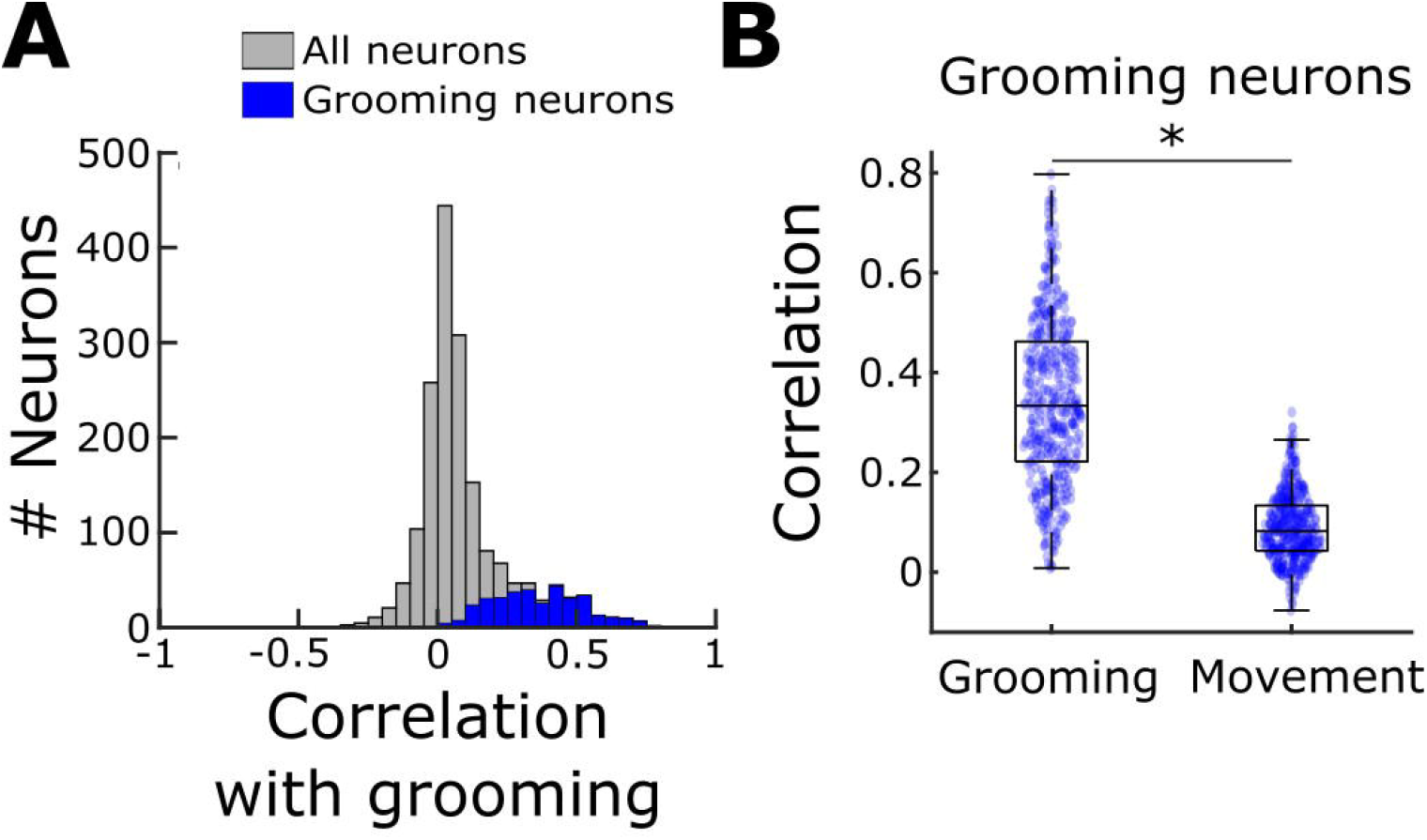
Correlation coefficients for all neurons across all mice with grooming behaviors and non-grooming forelimb movements. **A)** Pearson correlation coefficients for all neurons (gray) and the grooming neurons identified with Rastermap (blue) with grooming behaviors. All neurons, n=2198, and grooming neurons n=418 over 7 sessions across 4 mice, Wilcoxon rank-sum test, p<0.01. **B)** Pearson correlation coefficients for each of the grooming neurons within the population highlighted in A (blue), with grooming and non-grooming forelimb movements. n=356 neurons, paired t-test, p<0.05

